# Abdominal symptoms of invasive meningococcal disease are associated with the induction of plasminogen activator inhibitor in omental adipocytes

**DOI:** 10.1101/2025.02.26.640293

**Authors:** Muhamed-Kheir Taha, Damien Oliveira, Marion Gros, Myriam Aouiti-Trabelsi, Ala-Eddine Deghmane

**Affiliations:** Institut Pasteur, Université Paris Cité, Invasive bacterial infection. 28 ru du Dr Roux, 75015 Paris, France

**Keywords:** *Neisseria meningitidis*, abdominal manifestations, animal model, transcriptomic analysis, plasminogen inhibitor activator 1

## Abstract

Abdominal symptoms are increasingly reported in invasive meningococcal disease (IMD), but the underlying mechanisms remain unclear. We aimed to explore the pathophysiology of these presentations using an animal model. We utilized a collection of 20 meningococcal isolates that were either associated or not associated with abdominal presentations, which were injected intraperitoneally into transgenic mice expressing human transferrin. We employed histological examination, RNA sequencing (RNAseq) transcriptomic analysis, and reverse transcriptase real-time PCR to analyze tissue preparations of the mice’s omentum.

The 20 tested isolates induced similar levels of bacteremia in mice. However, isolates associated with abdominal presentations (mainly serogroup W isolates of clonal complex 11) caused thrombotic lesions in the blood vessels of the omentum, and they also induced a higher inflammatory response in the omentum with elevated levels of IL-6, TNF-alpha, and KC. Furthermore, these isolates induced higher expression of several genes, some of which are involved in coagulopathy, such as plasminogen activator inhibitor 1 (PAI-1). We also demonstrated that the PAI-1 encoding gene is overexpressed in adipocyte cells of the omentum. Lipopolysaccharide from the isolates associated with abdominal presentations, instead of whole bacteria, induced similar pathological findings.

During IMD, thrombosis formation in the omentum’s blood vessels is associated with a local induction of an inflammatory response and overexpression of the plasminogen activator inhibitor 1 encoding gene. These lesions can lead to thrombosis and hypoperfusion in the omentum, resulting in clinical abdominal presentations

**Author summary:** *Neisseria meningitidis*, commonly known as meningococci, is a bacterium that is transmitted directly from person to person through respiratory droplets. This bacterium causes invasive meningococcal disease (IMD), which can manifest in various symptoms beyond just meningitis. Notably, abdominal presentations, including abdominal pain and diarrhea, are increasingly being reported. The aim of our investigation was to elucidate the underlying mechanism of these abdominal symptoms.

To achieve this, we employed several experimental approaches and provided evidence that these symptoms are caused by the coagulation (clotting) of blood in the microvessels that surround the abdominal organs, such as the intestine. This clotting is triggered by the bacterium’s induction of a human enzyme that promotes coagulation. Notably, this is the first study to explore a mechanism underlying an extra-meningeal clinical form of *N. meningitidis* infection.

The enzyme responsible for this coagulation, plasminogen activator inhibitor 1, is a potential target for modulating the host response to IMD. Our findings have significant implications for the understanding of meningococcal pathophysiology and reveal additional potential targets for treatment.

## Introduction

*Neisseria meningitidis* (Nm), also known as meningococcus, is a Gram-negative, diplococcal bacterium found exclusively in humans. It is a commensal of the nasopharyngeal sphere. Transmission occurs through Human-to-human contact via respiratory droplets [1].

Acquisition of meningococcal isolates most often results in asymptomatic carriage, with a carriage prevalence of 10% in the general population [2]. However, this rate varies with age, being lower in infants, increasing during childhood, and peaking in adolescents and young adults [3]. Certain isolates of *N. meningitidis* can cross the respiratory epithelium and invade the bloodstream, causing septicemia that can progress to septic shock. Bacteria can cross the blood-brain barrier and invade the subarachnoid space in the central nervous system, causing cerebrospinal meningitis. Invasive meningococcal disease (IMD), even treated, leads to high mortality (10% of cases) and life-lasting sequelae (25% of survival cases) such as amputation, deafness, and neurological disorders in addition to an important cost of case management exceeding 11,000 euros per index case [4, 5].

Meningococcal invasiveness is determined by both host and bacterial factors, as well as viral co-infections [6]. Bacterial factors such as outer membrane proteins and the polysaccharide capsule contribute to meningococcal pathogenesis [7]. Capsular polysaccharides define twelve serogroups, six of which (A, B, C, Y, W, and X) are responsible for almost all IMD cases worldwide, although with a variable distribution [8]. Molecular typing methods such as Multi Locus Sequence Typing (MLST) and subsequently whole genome sequencing (WGS) have allowed the identification of hyperinvasive genotypes (also called hyperinvasive clonal complexes, CCs) among invasive isolates. The distribution of serogroups and clonal complexes varies temporally and geographically. For example, since 2011, there has been a global expansion of a serogroup W strain of the CC11, which seems to have originated in South America, then spread to the UK, and subsequently to the rest of Europe (this lineage is known as the South America-UK strain). Genomic analysis of CC11 isolates worldwide confirmed the emergence of the South America-UK strain [9]. Other lineages of serogroup W/CC11 have also shown worldwide spread, such as those linked to the previous emergence in 2000 of another strain named the Anglo-French-Hajj strain [9]. Isolates of the South America-UK strain seem to frequently cause atypical clinical presentations such as abdominal syndromes [10]. These abdominal presentations were hypothesized to be associated with the formation of microinfarcts in the blood vessels of the omentum, a highly vascularized fold of the peritoneum connecting the various abdominal viscera. Corroboratively, meningococci were detected in the blood vessels of the duodenum in a patient with IMD associated with abdominal pain [11]. Previously published genomic analyses have shown that isolates belonging to the South America-UK strain display allelic differences in genes involved in the biosynthesis of lipooligosaccharide (LOS) (the bacterial endotoxin) compared to isolates belonging to the Anglo-French-Hajj strain [10]. Meningococcal LOS is composed of three highly variable short oligosaccharide chains named alpha, beta, and gamma chains, whose presence and length depend on the expression of the phase-variable *lgt* genes [12]. Differences in the length and composition of the oligosaccharide chains and inner core dramatically alter the antigenic properties of LOS and form the basis of the classification into different immunotypes (L1 to L12)[13, 14].

This work aimed to mimic the pathophysiological process of these abdominal presentations in mice and test the role of meningococcal LOS in causing lesions that may lead to these abdominal symptoms.

## Results

### Characterization of selected isolates

To investigate the mechanism of abdominal presentation we initially selected a collection of isolates enriched in serogroup W, as this serogroup was reported to be more associated with abdominal presentations [10]. This collection consisted of 20 meningococcal isolates from various serogroups, which were notified (n=13) or not (n=7) to be associated with abdominal presentations at hospital admission. The isolates belonged to the following serogroups: B (n=3), C (n=2), Y(n=1), and W (n=14) (**Table 1**). Based on serogroups and clinical presentations, the isolates were categorized into four groups: W isolates with abdominal presentation (**Wabdo**), W isolates without abdominal presentation (**Wnonabdo**), non-W isolates with abdominal presentation (**NonWabdo**) and non-W isolates without abdominal presentation (**NonWnonabdo**) (**Table1**).

**Table 1.** Isolates used in the study and their characteristics.

| Id (PUBMLST) | Isolate | Relevant clinical form* | Site | Year | Serogroup | Clonal complex | *lgtA*‡ | *lgtG*‡ | Immunotype L3,7,9 |
| --- | --- | --- | --- | --- | --- | --- | --- | --- | --- |
| 41857 | 28585 | Abdo | Blood | 2016 | W | 11 | ON | OFF | + |
| 49253 | 28772 | Abdo | Blood | 2016 | W | 11 | ON | OFF | + |
| 51010 | 28840 | Abdo | Blood | 2016 | W | 11 | ON | OFF | + |
| 63155 | 31174 | Abdo | Blood | 2019 | W | 11 | ON | OFF | + |
| 85336 | 30634 | Abdo | Blood | 2019 | W | 11 | ON | OFF | + |
| 85884 | 30766 | Abdo | Blood | 2019 | W | 11 | ON | OFF | + |
| 88989 | 30823 | Abdo | Blood | 2019 | W | 11 | ON | OFF | + |
| 89221 | 30905 | Abdo | Blood | 2019 | W | 11 | ON | OFF | + |
| 89267 | 30924 | Abdo | Blood | 2019 | W | 11 | ON | OFF | + |
| 94153 | 31135 | Abdo | Blood | 2019 | W | 11 | ON | OFF | + |
| 84494 | 30434 | Abdo | Blood | 2019 | B | 32 | ON | OFF | + |
| 85524 | 30694 | Abdo | CSF | 2019 | B | 32 | ON | OFF | + |
| 85783 | 30714 | Abdo | Joint fluid | 2019 | C | 11 | ON | OFF | + |
| 53519 | 29116 | Non-abdo | CSF | 2017 | W | 11 | ON | ON | - |
| 53528 | 29133 | Non-abdo | CSF | 2017 | W | 11 | ON | ON | - |
| 57597 | 29590 | Non-abdo | Blood | 2018 | W | 11 | ON | ON | - |
| 61320 | 30105 | Non-abdo | Joint fluid | 2018 | W | 11 | ON | ON | - |
| 71393 | 31259 | Non-abdo | CSF | 2019 | B | 162 | ON | ON | - |
| 93476 | 31086 | Non-abdo | CSF | 2019 | C | 11 | ON | ON | - |
| 94137 | 31093 | Non-abdo | Blood | 2019 | Y | 167 | ON | ON | - |

\* Abdo indicates (association with abdominal presentation), Non-abdo indicates (no reported abdominal presentation).
‡ ON indicates the corresponding gens (*lgtA* or *lgtG*) is in-frame. OFF indicates the corresponding gens (*lgtA* or *lgtG*) is out of frame.
§ (+) and (-) indicates that the corresponding isolate reacts or does not react respectively with the L3,7,9 monoclonal antibody in ELISA.

The isolates were further characterized by whole genome sequencing, which allowed the extraction of clonal complexes (CCs) (**Table 1**). Most of the isolates (n=16) belonged to ST-11 (CC11) and were of serogroup W (n=14) or C (n=2) with the majority (n=11) associated with abdominal presentations, while five isolates were not (**Table 1**). The W/CC11 isolates, although highly related, clustered separately into two subgroups on the cgMLST network according to their association with abdominal presentation (**Fig.1**). Previous studies have suggested that LOS structure may differ between these two subgroups of W/CC11[10]. Therefore, we screened the WGS data to deduce the LOS immunotypes (named L1 to L12) in these 20 bacterial isolates [15]. This prediction was based on the role of different genes involved in LOS biosynthesis and the production of the enzymes encoded by these genes. The *lgtA* gene, which is involved in the biosynthesis of the alpha chain of LOS, was in the ON phase in the isolates associated with or without abdominal forms, predicting the presence of the L3,7,9 epitope. However, the *lgtG* gene, which is involved in the biosynthesis of the beta chain, was in the OFF phase, in the isolates associated with the abdominal forms, predicting the absence of this chain in these isolates but not in the isolates. Isolates not associated with abdominal forms showed the presence of the beta chain (**Table 1**). These data suggest that the alpha chain is present in all isolates, which should possess a complete lateral alpha chain and are predicted to react with the monoclonal antibody L3,7,9.

**Figure 1.**
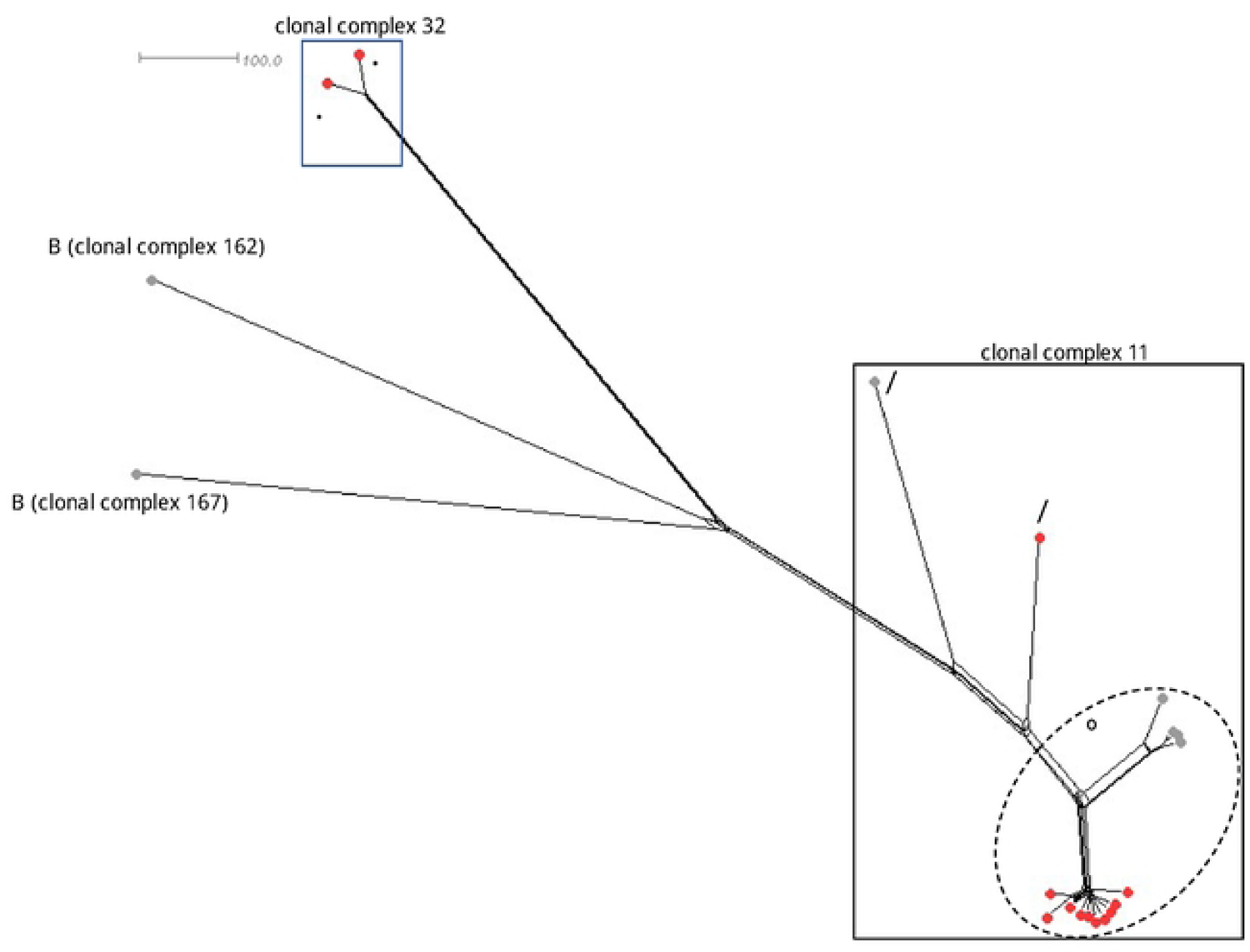
A neighbor-net network of the 20 isolates used in this study according to their core-genome MLST. Serogroups and clonal complexes are indicated. Individual isolates are represented by colored circles. Red circles denote isolates associated with abdominal presentations, while grey circles correspond to isolates that were not reported to be associated with abdominal presentations.

Using whole bacteria ELISA, we detected the presence of the L3,7,9 epitope, as predicted by the sequence analysis. All of the isolates associated with abdominal forms presented detectible L3,7,9 epitopes. Isolates not associated with abdominal forms did not show detectable L3,7,9 epitopes, despite the sequence prediction. This lack of recognition is associated with the presence of the beta chain (*lgtG* ON gene). This suggests that the beta chain may mask the accessibility of the L3,7,9 epitope on the alpha chain of LOS.

### Meningococcal isolates associated with abdominal syndromes induce micro-infarcts in the omentum

We used experimental infections in mice to characterize the lesions caused by the 20 selected isolates and to assess their virulence. Transgenic mice expressing human transferrin were inoculated intraperitoneally with 0.5 mL of a bacterial suspension containing 5 × 10^7^ colony-forming units, (CFU) per mL of the isolates. Control mice received 0.5 mL of buffer alone intraperitoneally. The clinical status of the mice was evaluated six hours post-infection using a combined scoring system based on fur appearance, strength, and peripheral temperature. Infected mice exhibited a clinical decline compared to non-infected controls.

Bacteremia levels were determined six hours post-infection, and mean values were similar across all infected groups (**Fig. 2A**). Similar bacteremia levels were observed when mice were grouped based on infection with isolates associated or not associated with abdominal presentations, regardless of serogroup.

**Figure 2.**
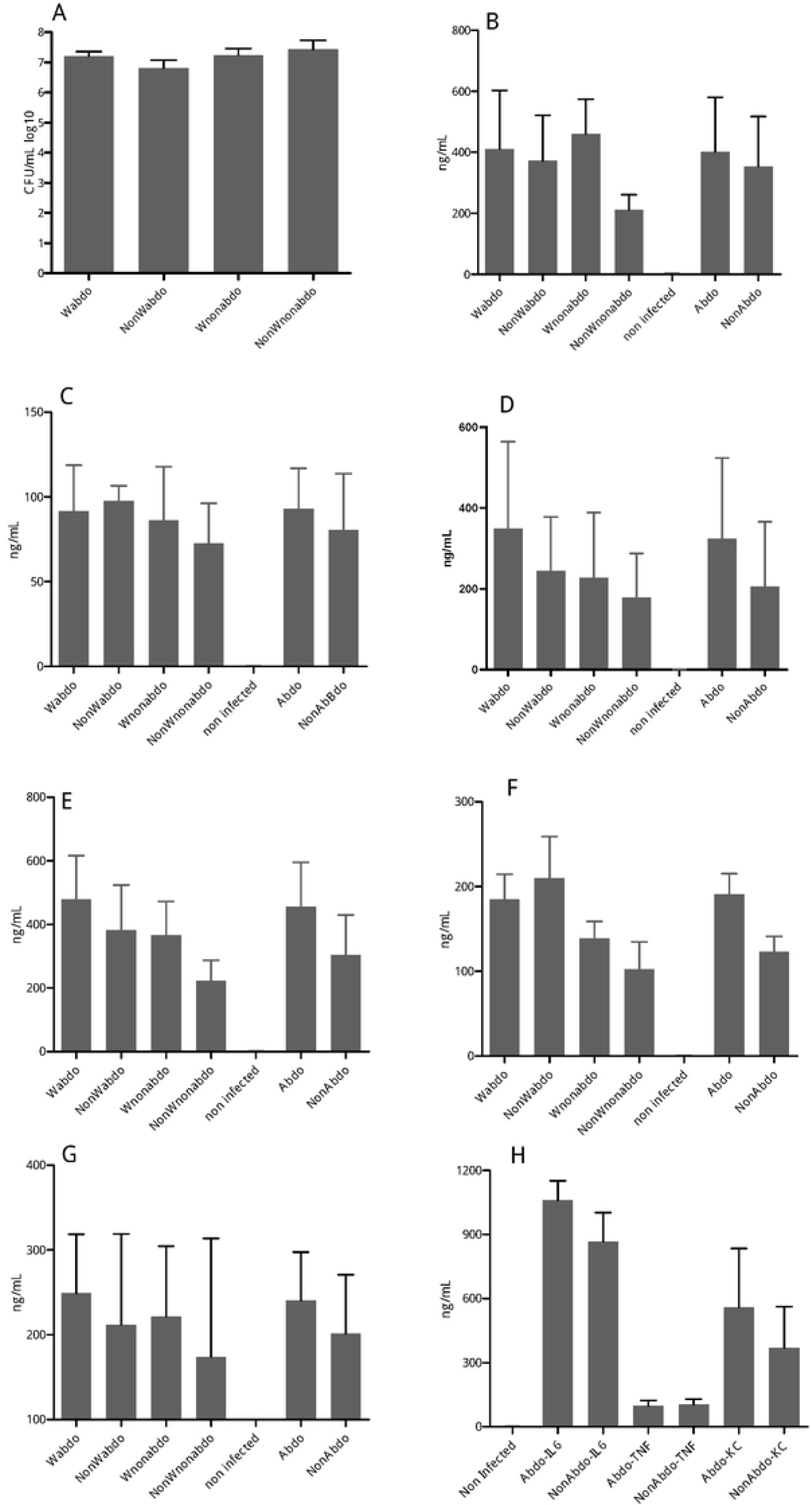
Bacteremia and cytokine production in transgenic mice expressing the human transferrin and infected by intra-peritoneal injection of 5 x 10^7^ CFU/mL of the 20 isolates used in this study. The groups of mice are indicated under the X-axis: W isolates with abdominal presentation (**Wabdo**), W isolates without abdominal presentation (**Wnonabdo**), non-W isolates with abdominal presentation (**NonWabdo**), non-W isolates without abdominal presentation (**NonWnonabdo**), all isolates with abdominal presentations (**Abdo**), and all isolates without abdominal presentations (**NonAbdo**). (A) Levels of bacteremia in mice expressed as log10 of CFU/mL for each group of mice as defined above. (B) Levels of IL-6 in the omentum from mice were expressed as ng/mL for each group of mice as defined above. (C) Levels of TNF-alpha in the omentum from mice were expressed as ng/mL for each group of mice. (D) Levels of KC in the omentum from mice were expressed as ng/mL for each group of mice. (E) Levels of IL-6 in the sera from mice were expressed as ng/mL for each group of mice. (F) Levels of TNF-alpha in the sera from mice were expressed as ng/mL for each group of mice. (G) Levels of KC in the sera from mice expressed as ng//mL for each group of mice. (H) Levels of cytokines were expressed as ng/mL in the omentum from mice injected by intraperitoneally with LOS (groups of mice are indicated for the three tested cytokines IL-6, TNF-alpha, and KC). Data are expressed as the mean concentration for each tested cytokine. Non-infected mice were used as controls.

Cytokine levels in blood and omentum samples were measured. IL-6, KC, and TNF-α levels were similar in both sites in infected mice. However, mice infected with Wabdo isolates showed higher cytokine levels, while those infected with NonWnonabdo isolates showed lower levels, although these differences were not statistically significant (**Fig. 2B-G**).

Given that our data suggested potential differences in lipopolysaccharide (LOS) between isolates associated or not with abdominal presentations, we injected purified LOS from representative isolates of both types. IL-6, KC, and TNF-α were measured six hours post-injection in both blood and omentum samples. While cytokines were not detectable in serum samples, they were detected in omentum fractions, with higher levels of IL-6, TNF-α, and KC in mice injected with LOS from isolates associated with abdominal syndromes. However, these differences did not reach statistical significance (**Fig. 2H**). Histological analysis of 5 μm omentum sections stained with eosin and hematoxylin (HE) revealed that the overall omentum structure was conserved in mice infected with both Wabdo and Wnonabdo isolates compared to non-infected mice. However, hemorrhagic spots and deformed vessels with accumulating fibrin were observed in sections from Wabdo isolate-infected mice, suggesting coagulation lesions in the omentum (**Fig. 3**).

**Figure 3.**
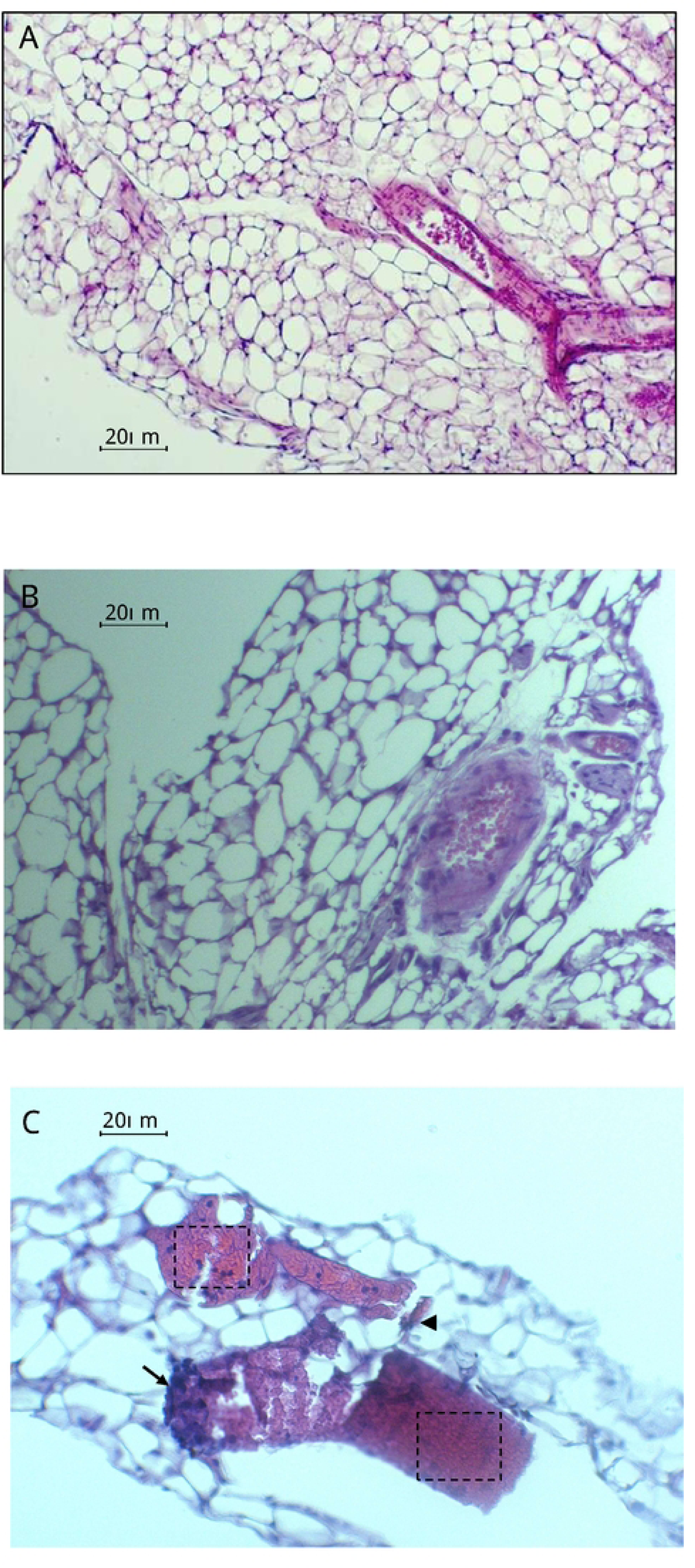
Hematoxylin and eosin (HE) staining of omentum sections (X20 magnification) from an uninfected mouse (A), a mouse infected by a Wnonabdo isolate (B), or a Wabdo isolate (C). Thrombotic lesions with red blood cells located in the center and surrounded by fibrin accumulations (dotted squares) are observed. Red blood cells are alsoobserved around a vessel (arrowhead) with an endothelialization on the surface blood clot (arrow). Size bars are indicated.

### Transcriptomic analysis of the omentum

RNAseq experiments were conducted on omentum fractions extracted from mice six hours post-infection with isolates associated or not with abdominal presentations. Differentially expressed genes compared to non-infected mice are listed in Supplementary Table 1. Genes expressed in both conditions were involved in inflammatory responses, such as serum amyloid A1 and A2 encoding genes (*saa1* and *saa2*).

We focused on differentially expressed genes in the omentum of mice infected with isolates associated with abdominal presentations. Several genes involved in coagulation homeostasis, such as the ceruloplasmin encoding gene (*cp*) and genes of the serpin family, were identified. Notably, the gene encoding serpin e1 (plasminogen activator inhibitor 1, *pai-1*) was upregulated, suggesting procoagulative activity and corroborating the observation of micro-infarcts in omentum sections (**Fig. 3** and **Supplementary Table 1**).

Specific genes were further tested by RT-rtPCR using primers and probes for several genes involved in the coagulation cascade (**Table 2**). Total RNA was obtained from omentum preparations of mice infected with isolates associated or not associated with abdominal presentations. Targeted gene expressions were compared between these two groups using the actin-encoding gene as an internal reference and non-infected mice as a background control. Genes encoding IL-6, TNF-α, serpin e1 (*pai-1*), serpin b2 (*pai-2*), and factor 3 (tissue factor of the coagulation cascade) were hyperexpressed in the omentum of mice infected with isolates associated with abdominal presentations compared to those infected with isolates not associated with abdominal presentations. Notably, the PAI-1-encoding gene showed significant overexpression (**Fig. 4**), corroborating the RNAseq results.

**Figure 4.**
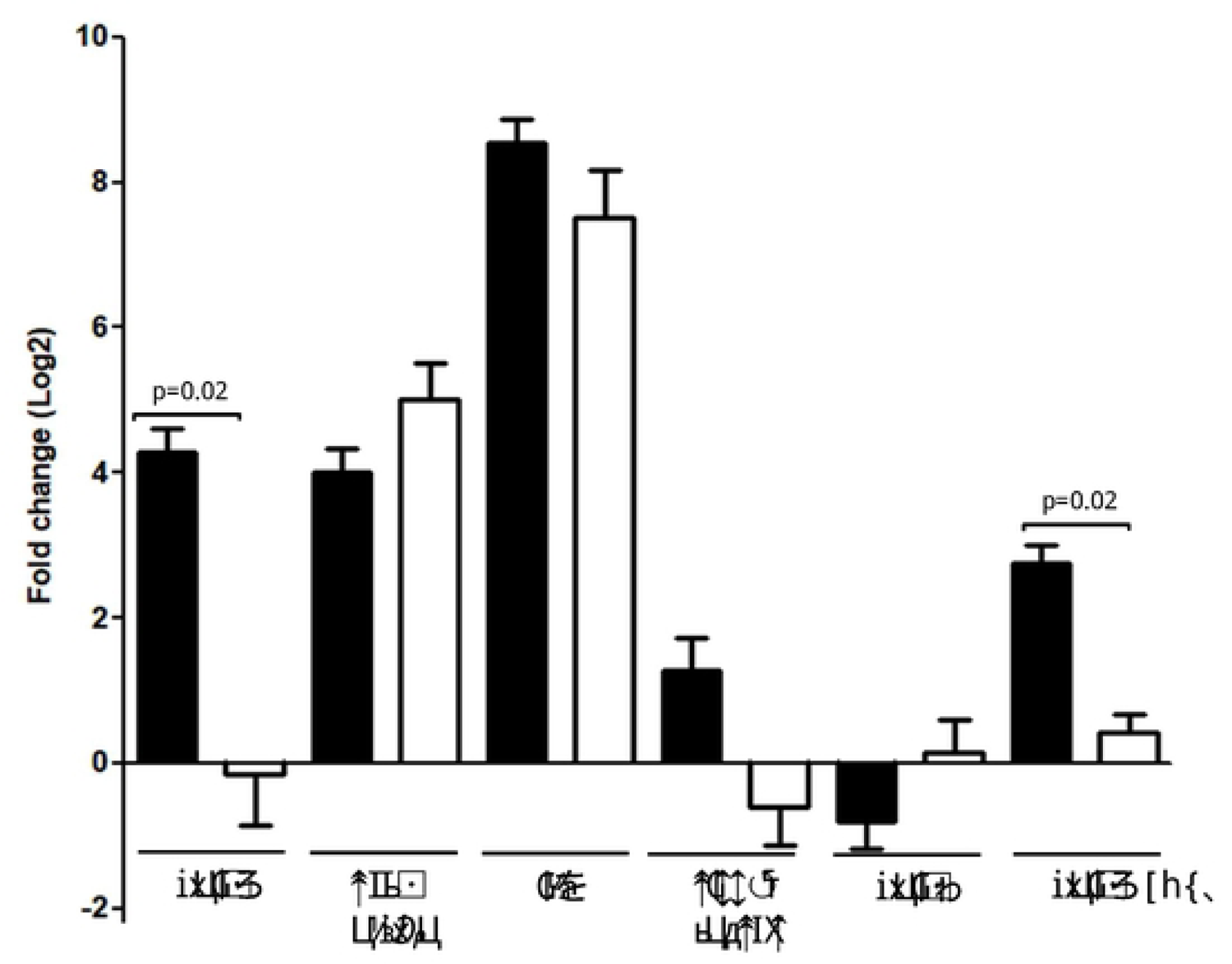
Comparison of genes differentially expressed in the omentum during infection. Expression of selected genes in omentum from transgenic mice expressing the human transferrin at 6h post-intraperitoneal infection by isolates associated (black) or non-associated (white) with abdominal presentations. Expression was quantified by reverse transcription-quantitative PCR (RT-qPCR). The gene encoding beta-actin was used as an endogenous housekeeping gene. Non-infected mice were used as a reference. Each bar represents the mean of at least three infections (*p*-value is indicated for significant differences).

**Table 2.**
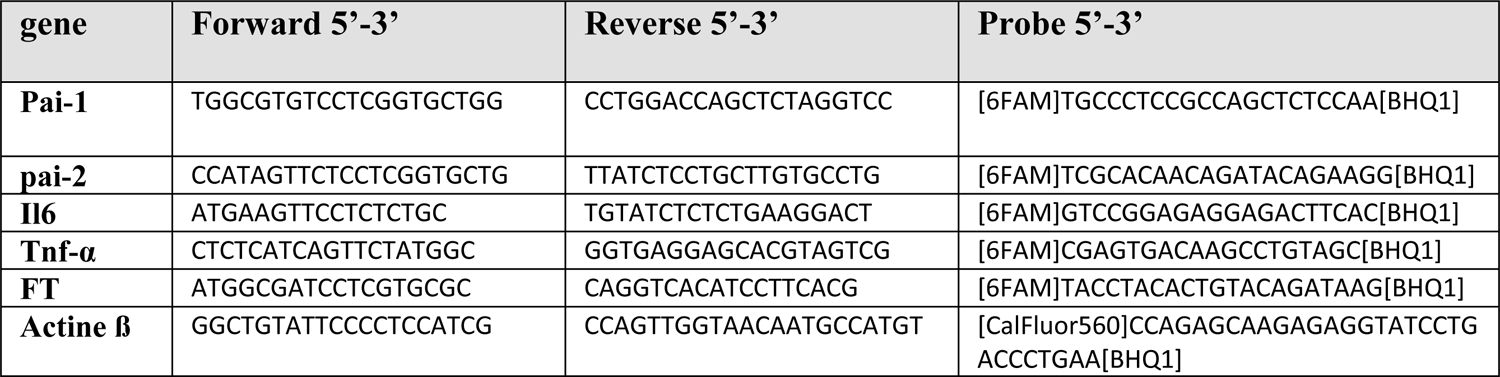
Primers and probes.

Similar RT-rtPCR analysis was performed on RNA extracted from mice injected with purified LOS from representative isolates of both types. Significant induction of the PAI-1-encoding gene was observed in mice injected with LOS from isolates associated with abdominal presentations (**Fig. 4**).

### RT-rtPCR in situ

To identify the cells involved in the overproduction of the PAI-1 encoding gene in the omentum, we performed RT-rtPCR on histological sections of the omentum using 5 μm sections. The data (**Fig. 5**) showed that sections from mice infected with viable bacteria (Wabdo) or with purified LOS from the same isolate exhibited staining in the adipocytes of the omentum compared to sections from non-infected mice. This suggests that the PAI-1 encoding gene is overexpressed in adipocyte cells, which appear to be a major cell type responding to meningococcal infection or LOS treatment.

**Figure 5.**
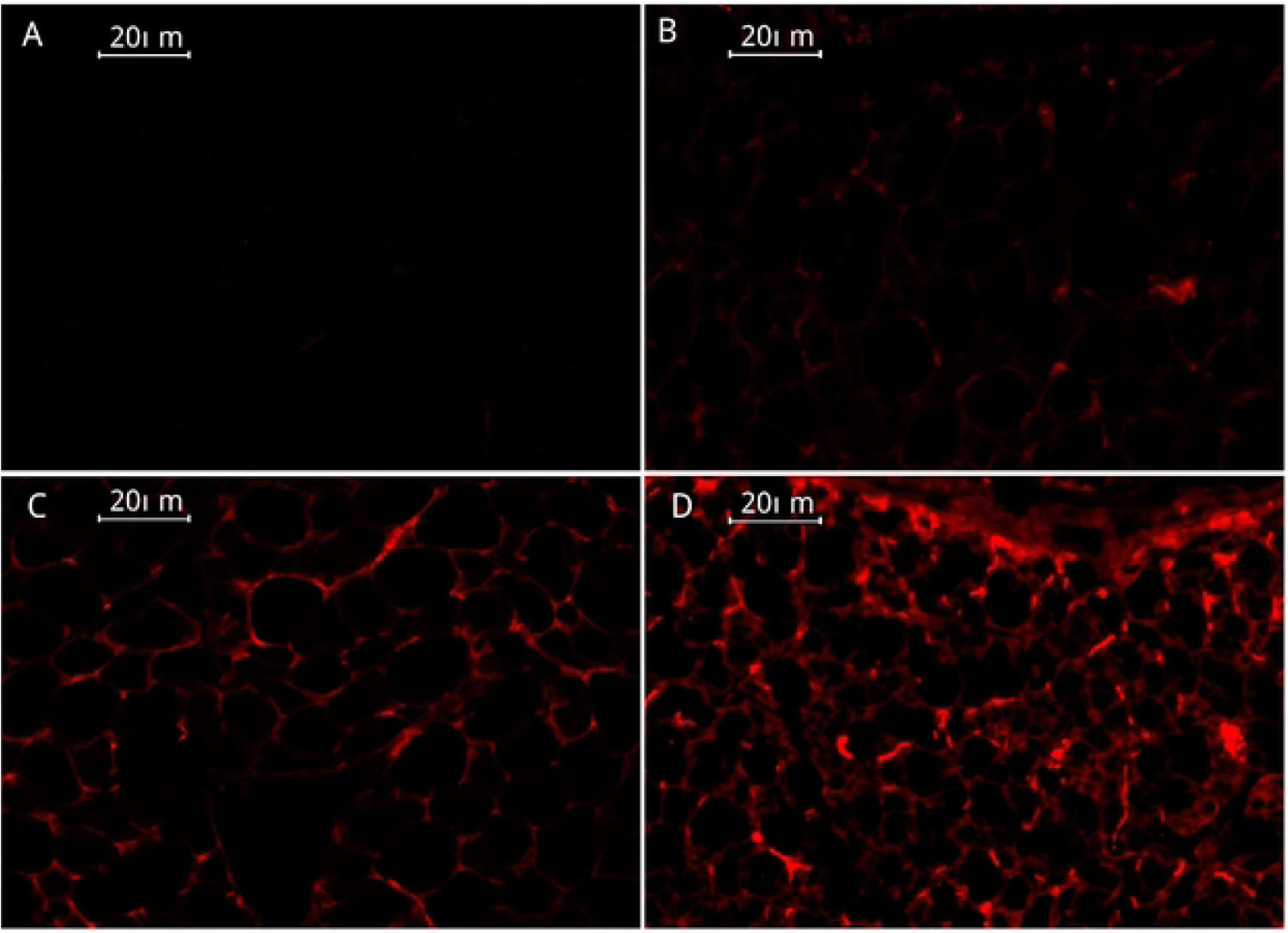
In situ reverse transcription-PCR (RT-PCR) analysis of plasminogen activator-inhibitor-1 (*pai-1*) mRNA in omentum sections. Several 5µm sections of the omentum were deparaffinized and then prepared as described in the Materials and Methods section. In situ RT-PCR reactions were performed using the SuperScript IV VILO Master Mix kit (Thermo Fisher Scientific, Illkirch, France). Primers and probe to detect the expression of the *pai-1* gene (**Table 2**) were used. Sections were examined using a fluorescent microscope Zeiss Axio Imager D1 coupled to AxioCam MRm vers.3 (Carl Zeiss, Germany). *pai-1* transcripts are labeled in red. (A) An unlabeled section, (B) Section from a non-infected mouse, (C) Section from a mouse infected by a Wabdo isolate, and (D) Section from a mouse injected with LOS from the Wabdo isolate in (C). Size bars are indicated.

## Discussion

IMD is often synonymous with meningitis; but, a variety of other clinical presentations have been increasingly reported, especially since the relaxation of COVID-19 control measures. These non-meningeal forms primarily include abdominal presentations such as abdominal pain and diarrhea [16]. Our study suggests that meningococcal isolates associated with abdominal forms induce higher local inflammatory reactions than those not associated with abdominal forms. Meningococcal LOS appears to be involved in the induction of the inflammatory response, as the injection of purified LOS from corresponding isolates correlated with the induction of a local inflammatory response. This response mimicked the inflammatory response observed with whole bacteria and seems correlated with the presence of the L3,7,9 epitope. The L3,7,9 immunotype, corresponding to the alpha chain of LOS, is associated with virulence and significant induction of inflammatory cytokines TNF-α and IL-6. In contrast, isolates with different immunotypes or truncated alpha chains (L8 immunotype) do not induce such responses [17]. Isolates of serogroup W clonal complex 11 (W/CC11), which harbor the L3,7,9 immunotype, seem to induce a stronger inflammatory response in the omentum. The non-accessibility of this epitope appears to reduce the inflammatory response. Specifically, if the *lgtG* gene is in the ON configuration, the beta chain is predicted to be present and can mask the alpha chain containing the L3,7,9 epitope. This may explain the lower levels of IL-6, KC, and TNF-alpha after 6 hours of injection with whole bacteria and purified LOS from isolates not associated with abdominal presentations, compared to those associated with abdominal presentations. Our study also suggests that other bacterial factors may be involved, as the responses observed with whole bacteria were systemic and stronger, while LOS injection provoked only local responses.

Our data also suggest that infection with whole bacteria or injection with LOS promotes coagulation and thrombus formation. This effect seems to result from a differential regulation of host genes involved in fibrinolysis, such as the PAI-1 encoding gene. Fibrinolysis is the process of degrading fibrin clots, beginning with the conversion of plasminogen into plasmin, which hydrolyzes fibrin clots. Counter-regulatory mechanisms exist to prevent excessive activation of fibrinolysis by inhibitors like PAI-1 and PAI-2. During IMD, LOS induces the release of PAI-1 by monocytes, leading to a reduction in fibrinolysis and contributing to thrombotic lesions [18, 19]. Our study also supports the findings that high production of PAI-1 is reported in IMD and high production of PAI-1 were reported in IMD [20].

Regarding the inflammatory response, isolates associated with abdominal presentations (Wabdo) and their corresponding LOSs induced stronger expression of the PAI-1 gene compared to other isolates. Upon induction, plasminogen activator inhibitor represses plasmin production, reduces fibrinolysis, and promotes coagulation [21]. This is consistent with the vascular lesions observed in the omentum of mice infected by Wabdo isolates or treated with their LOS. These results correlate with studies in patients with IMD, showing that individuals with severe IMD (e.g., purpura fulminans characterized by disseminated intravascular coagulation lesions) have high levels of endotoxin and PAI-1, establishing a procoagulant state compared to patients with milder forms of IMD [22]. Additionally, N. meningitidis endotoxin (LOS) induces the release of PAI-1 by monocytes [19].

Furthermore, a common functional deletion polymorphism in the promoter region of the *pai-1* gene leading to the (4G/5G) genotype has been reported. Subjects carrying this polymorphism produce higher concentrations of PAI-1, develop more severe coagulopathy, and are at greater risk of death during meningococcal sepsis [20]. Besides, its role in the regulation of fibrinolysis, PAI-1 has also been found to be associated with insulin resistance, and diabetes mellitus [23, 24]. Hence, increased PAI-1 levels may play a critical role in the development of impaired insulin sensitivity during meningococcal sepsis. This aligns with the frequent and significant hyperglycemic episodes associated with insulin resistance during acute and severe meningococcal sepsis [20, 25].

Our data suggest that adipocytes in the omentum may serve as a source of PAI-1, contributing to elevated plasma PAI-1 levels observed during meningococcal infection. The link between adipocytes and proinflammatory balance, the coagulation pathway, and impaired endothelial function is well-established in several metabolic disorders like atherosclerosis [26]. Our data also substantiate the link between these pathophysiological factors and infectious diseases [27]. Our study has several limitations. The association of an isolate with an abdominal presentation depends on the patient’s history and may be overlooked if not specifically solicited [10]. However, our study underscores the need to include IMD in the differential diagnosis of abdominal syndromes associated with systemic symptoms and signs. Another limitation is the difficulty of extrapolating results obtained in mice to humans. The genes encoding the serpin family may differ in their function and/or regulation between humans and rodents [28]. Finally, the meningococcal tropism for the blood vessels of the omentum remains to be elucidated.

In conclusion, abdominal presentations of IMD may result from mesenteric hypoperfusion related to thrombosis formation in mesenteric vessels. Our study supports the hypothesis that a local induction of an inflammatory response leads to thrombotic lesions, which seem to result from the specific action of PAI-1 on neighboring blood vessels.

## Material and Methods

### Characterization of meningococcal isolates

We selected 20 representative isolates from the National Reference Center for meningococci and *Haemophilus influenzae* (NRCMHi) database that were associated or not with reported abdominal manifestations. The isolates were characterized phenotypically (capsule expression) and by whole genome sequencing as previously reported [29] and their characteristics are depicted in **Table 1**. The genomic data were used to extract sequences of genes involved in LOS biosynthesis using the tool available on https://pubmlst.org [30]. LOS is composed of a lipid part (lipid A) and an oligosaccharide part that contains two conserved KDO molecules (I and II, keto-D-manno-octulosonic acid) and two conserved heptose molecules (Hep I and II). Unlike enterobacteria, meningococcal LOS does not contain highly repeated lateral saccharide chains. It is composed of a chain alpha (connected to Hep I), a chain beta, and a chain gamma (connected to Hep II). The genes involved in the biosynthesis of LOS are *lgtA*, *lgtB*, *lgtC*, *lgtD*, *lgtE*, *lgtF*, *lgtG* and *lgtH*, with several of them that undergo phase variations ON or OFF and lead to alternative LOS structures that are defined as immunotypes L1 to L12 [15, 31].

### Purification of LOS

LOS purification was performed from isolates cultured at 37°C under 5% CO2 on GC medium (Thermo Fisher Scientific, Illkirch, France), supplemented with Kellog supplements I and II for 18-20h [32]. Bacterial suspensions were prepared by harvesting 2 plates in physiological saline and treated in acid phenol (vol/vol) precipitated with acetone. The pellets containing the crude LOS were subjected to dialysis overnight against 1 L of bi-distilled water and then LOS were recovered by several cycles of ultracentrifugation at 100,000xg as previously described [33]. LOS preparations were used after standardization to concentrations equivalent to the bacterial suspension concentrations used in infection (5E7 CFU/mL). Whole-bacteria ELISA detection of the LOS epitope 3,7,9 was performed using the specific monoclonal antibody to this epitope [34, 35].

### Mice infection

Transgenic mice expressing the human transferrin were previously described by our laboratory [36]. Human transferrin in these transgenic mice provides the required iron for bacterial growth as *N. meningitidis* utilizes specifically human iron sources such as the human transferrin [37]. We used 8-12 weeks-old female mice for infection. Mice were infected by intraperitoneal injection of 0.5 mL of a bacterial suspension of 5x 10^7^ CFU/mL. Infection was also performed with purified LOS corresponding to viable bacterial infectious dose (5×10^7^ CFU/mL). Clinical score, composed of a combined score on fur, strength, and peripheral temperature of mice, was measured before and 6 h after infection as previously described [38].

Bacterial loads at 6 h post-infection were determined by plating serial dilutions of blood on GC medium plates. Aliquots of blood were also used for cytokines assays. After 6h of infection, mice were euthanized by an intramuscular injection of an overdose of a mixture of 10 mg/kg xylazine (Bayer, Puteaux, France) and 5 mg/kg ketamine (Merial, Lyon, France).

### Omentum preparation

A laparotomy was performed in euthanized mice. A syringe containing approximately 5mL of air, mounted on a 25G-needle, was then implanted at the colon level allowing the swelling of the intestine and the detachment of the omentum. Two omentum samples were taken on PureLink™ bead tubes (Thermo Fisher Scientific, Illkirch, France) and stored at −80°C. One sample was used for total RNA extraction using the PureLink® RNA mini kit (Thermo Fisher Scientific, Illkirch, France) according to the instruction of the manufacturer. The yield and purity of the RNAs were determined with the NanoDrop® One spectrophotometer (Thermo Fisher Scientific, Illkirch, France) according to absorbance measurements at 260 nm, 280 nm and 230 nm.

The second sample was used for cytokine assays. Proteins were prepared after grinding and centrifugation. The quantity/concentration of proteins was measured with the NanoDrop® One spectrophotometer (Thermo Fisher Scientific, Illkirch, France) and standardized across all samples to 1mg/L. Quantification of cytokines (IL-6, TNF-α and KC) in the blood and in the omentum of mice was carried out by ELISA method using DuoSet® kit (R&D systems, Noyal Châtillon sur Seiche, France) following the instructions of the manufacturer.

A piece of the omentum was also cut with scissors, recovered on biopsy foam, and immediately immersed in 10% formalin for at least 24 hours. It was then fixed in 70% alcohol. Samples were then dehydrated overnight, embedded and cut in paraffin at 5 µm horizontal sections before staining with hematoxylin and eosin (HE) as previously described [38].

### Transcriptional analysis

Transcriptomic analysis by RNAseq with the Illumina HiSeq2500®146 genomic sequencer was performed using total RNA prepared from omentum of mice infected with live bacterial isolates (associated or not with an abdominal syndrome) or non-infected. We analyzed mice that were infected by isolates that were associated with abdominal manifestations (n=3), and isolates that were not associated with abdominal manifestations (n=3). Non-infected mice were also used. This analysis made it possible to detect genes differentially expressed in the omentum of mice infected with isolates associated with abdominal syndromes during infection in humans. The RNAseq reads are aligned to the reference genome (*Mus musculus*) using Bowtie-generating genome/transcriptome alignments. TopHat identifies the potential exon-exon splice junctions of the initial alignment [39]. Then Cfflinks identifies and quantifies the transcripts from the preprocessed RNA-Seq alignment assembly (fragment per kilobase per million mapped reads, FPKM) [39, 40]. After this, Cuffmerge merges the identified transcript pieces into full-length transcripts and annotates the transcripts. Finally, merged transcripts from two or more samples /conditions are compared using Cuffdiff to determine the differential expression levels at the transcript and gene level including a measure of significance between samples/conditions [39, 40]. Differentially expressed genes were defined in comparison with non-infected mice. A significantly differentially expressed gene was defined if present in the three isolates in each of the two conditions (association or not with abdominal presentations) [41]. Data of the RNAseq are available on the BioSample database of the NCBI with the accession numbers SAMN41039157, SAMN41039158, SAMN41039159, SAMN41039160, SAMN41039161, SAMN41039162, SAMN41039163.

Additionally, reverse transcription real-time PCR (RT-rtPCR) was also performed for several targeted genes: plasminogen activator inhibitor-1 and 2 encoding genes (*pai-1* and *pai-2*), TNF-alpha encoding gene, IL6 encoding gene, tissue factor encoding gene, and beta-actin encoding gene. The primers and Taq Man probes characteristics of the targeted genes identified by the above analysis of FPKM are listed <u>in</u> **Table 2**. We used SuperScriptIV VILO™ Master Mix kit (Thermo Fisher Scientific, Illkirch, France) according to the instruction of the manufacturer with 2.5 μg of RNA. The obtained complementary DNA (cDNA) was stored at −80°C until use. cDNA was used in real-time PCR that was performed in a final volume of 25 μL with an AB 7300 Real-Time PCR system (Applied Biosystems). To quantify the expression of targeted genes, the relative ΔΔCt method was used by calculating the differences in scores between treated samples (infected mice) and untreated samples (non-infected mice) using the actin-encoding gene as an internal reference and using non-infected mice for the background. These scores correspond to differences in cycle thresholds between the gene of interest and a housekeeping gene (actin-encoding gene) [42].

Reverse-Transcriptase-real time PCR (RT-rtPCR) in situ was used to determine the tissue expression of the serpin (**ser**in **p**rotease **in**hibitor) encoding genes. Histological sections of 5 μm thickness were deparaffinized through two xylene washes for 5 minutes each, then rehydrated by 5 minutes washing through several ethanol baths at decreasing concentrations. Finally, the slide was washed for 5 minutes in two PBS successive baths and then in two bi-distilled water baths. A fixation step with 4% paraformaldehyde at 4°C for 1 hour was performed. Subsequently, cells were lysed on the slide for 10 minutes using the Cells-To-Ct kit according to the manufacturer’s instructions (Thermo Fisher Scientific, Illkirch, France). Reverse transcription was then performed for the targeted gene using SuperScriptIV VILO™ Master Mix kit (Thermo Fisher Scientific, Illkirch, France) according to the manufacturer’s instructions. The sections were covered with a coverslip and placed in the thermocycler. The thermocycler program consists of a first reverse transcription step (50°C, 5 min; 1 cycle), a second one that allows the inactivation of the RT and the initial denaturation (95°C, 20 sec; 1 cycle), and an amplification (95°C 15 s, 60°C 1min; 40 cycles). The slides were examined using a fluorescent microscope Zeiss Axio Imager D1 coupled to AxioCam MRm ver.3 (Carl Zeiss, Germany). Digital images were acquired using appropriate filters and combined using the Axiovision Rel. 4.6 software (Carl Zeiss).

### Statistical analysis

Statistical analyses were performed using Prism 9.1.0 software (GraphPad Inc. San Diego, CA, USA). *P* values <0.05 were considered statistically significant. Student t-test and non-parametric tests for quantitative variables were used to compare at least two independent experiments.

